# Interpretable Deep Learning of Endogenous Fluorophore Concentration and Entropy for Lung Tissue Classification

**DOI:** 10.64898/2026.10.26.762057

**Authors:** Zhuo Yin, Yunsong Guo, Leslie Virany, Min Xu

## Abstract

Fluorophore concentration measures and local histogram entropy describe different aspects of endogenous fluorophore maps. We investigate whether their combination improves discrimination between normal and cancerous lung tissue and which features contribute to predictions. Chemometric fluorescence microscopy (CFM) provides concentration maps of tryptophan, elastin, reduced nicotinamide adenine dinucleotide (NADH), and flavin adenine dinucleotide (FAD). A neural network based on generalized additive models with structured interactions (GAMI-Net) is used to compare concentration-only, entropy-only, and combined representations through explicit main effects and pairwise interactions. The mean areas under the receiver operating characteristic curve (ROC AUCs) across five-fold cross-validation were 95.9% ± 0.7%, 96.4% ± 1.6%, and 98.4% ± 0.7%, respectively; variability is reported as standard deviation (SD). Their aggregate differences are consistent with the hypothesis of complementary predictive information. FAD entropy, NADH concentration, and NADH entropy were the leading main effects in the combined model. The learned log-odds contributions include a U-shaped elastin-entropy relationship and nonmonotonic tryptophan relationships. By exposing how concentration and entropy enter predictions, this interpretable framework turns label-free fluorescence classification into specific, testable hypotheses about tissue metabolism and extracellular-matrix organization, motivating specimen-grouped and external validation.

## 1. Introduction

Carcinogenesis can alter tissue biochemistry and organization. Lung cancer caused approximately 1.8 million deaths worldwide in 2020 and was the leading cause of cancer mortality [1]. Earlier detection and accurate diagnosis remain central to reducing this burden [2]. Endogenous fluorescence provides biochemical and structural contrast in unstained tissue and offers a basis for quantitative optical assessment [3, 4].

The shift toward aerobic glycolysis associated with the Warburg effect changes energy requirements and redox regulation [5, 6]. NADH and FAD are fluorescent metabolic cofactors whose signals reflect aspects of cellular redox state; their relative fluorescence has been used to characterize metabolic differences in cancer models [4, 7, 8]. Elastin provides structural contrast from the extracellular matrix, whose remodeling accompanies tumor progression [9, 10]. Tryptophan metabolism links cellular nutrient use to immune regulation, including immunosuppressive catabolic pathways [11, 12]. Together, these fluorophores probe complementary aspects of tissue composition and function. Their spatial heterogeneity may provide information beyond concentration alone.

Quantitative chemometric wide-field fluorescence microscopy (CFM) combines broadband imaging, non-negative matrix factorization (NMF), and reference calibration to obtain concentration maps of endogenous fluorophores [13]. Fluorophore concentration describes abundance but does not fully characterize the spatial distribution in its map. In particular, different distributions can have the same mean concentration. Local histogram entropy summarizes the frequencies of pixel values within a neighborhood and may therefore provide predictive information beyond abundance. Higher entropy indicates a greater diversity of pixel values within the neighborhood. However, rearranging values within a neighborhood preserves its histogram and entropy, so entropy does not uniquely describe spatial arrangement or tissue architecture.

Xu et al. (2022) reported that combining concentration and entropy improved support vector machine (SVM) classification relative to concentration alone for lung-tissue classes (normal, perilesional, and cancerous) and for cancer grades 1–3 [14]. This prior evidence motivates evaluating the two feature groups together; their predictive contribution in the present binary normal-versus-cancerous task must still be tested. Our contribution is to examine their discrimination and learned relationships within an interpretable model, rather than to introduce the combination of concentration and entropy.

Machine learning has expanded the capabilities of optical microscopy through image analysis, virtual staining, and tissue classification [15–18]. However, predictive performance alone does not explain how individual biochemical features contribute to a decision. SVMs and convolutional neural networks (CNNs) can use combined features and can be interpreted, but their standard formulations do not provide the same explicit additive decomposition into individual-feature and pairwise functions. GAMI-Net is a neural network based on generalized additive models with structured interactions [19]. Its main-effect curves and interaction surfaces expose how concentration and entropy enter predictions while allowing nonlinear relationships. For binary classification, the intercept and these functions sum to the model’s log odds that a sample is cancerous rather than normal tissue; the logistic function converts this sum to a predicted probability.

We hypothesize that combining fluorophore concentration and entropy improves discrimination relative to either feature group alone. We test this hypothesis by comparing concentration-only, entropy-only, and combined GAMI-Net models. We then use the models’ main effects and pairwise interactions to identify the contributing features and feature pairs and to examine their functional relationships and biological plausibility. Section 2 describes the feature construction, model, and validation protocol; Section 3 reports classification performance and model explanations; and Section 4 considers biological interpretation and limitations.

## 2. Methods

Figure 1 illustrates how the mean concentration and mean local entropy of each fluorophore are extracted from nonoverlapping sub-images. Three feature groups are evaluated: concentration only (four concentrations), entropy only (four entropies), and concentration and entropy combined (all eight features). GAMI-Net decomposes the classification log odds into main effects and pairwise interactions, so the feature-group comparison is followed by an examination of how the measured features contribute to predictions.

**Fig. 1.**
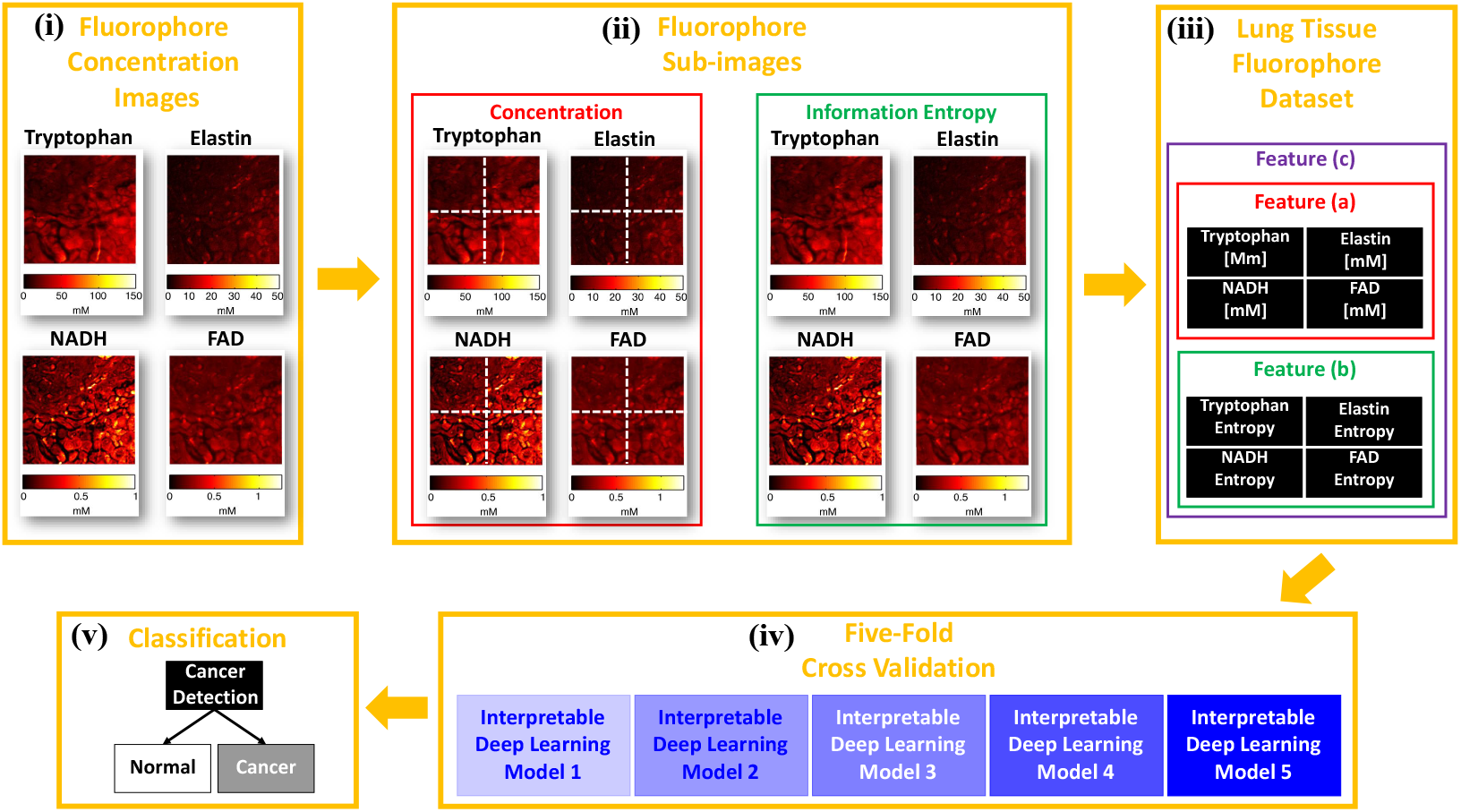
Workflow from concentration maps to interpretable classification. (i) Maps are partitioned into nonoverlapping sub-images. (ii) Local entropy maps are computed from the concentration sub-images. (iii) Sub-image mean concentrations and mean entropies provide eight features. (iv–v) GAMI-Net is trained and evaluated with five-fold cross-validation for concentration-only, entropy-only, and combined inputs. The partitioning and reporting conventions are specified in Sections 2.5.1 and 2.6.

### 2.1. Epi-fluorescence experiment and tissue images

An unstained, formalin-fixed, paraffin-embedded lung tissue microarray (BioMax, USA), comprising normal (*n* = 36) and cancerous (*n* = 52) specimens from a cohort of 18 patients, was imaged using an epifluorescence microscope (Axio 40cfl, Zeiss) [13]. Figure 2 illustrates the shared excitation and collection optics.

**Fig. 2.**
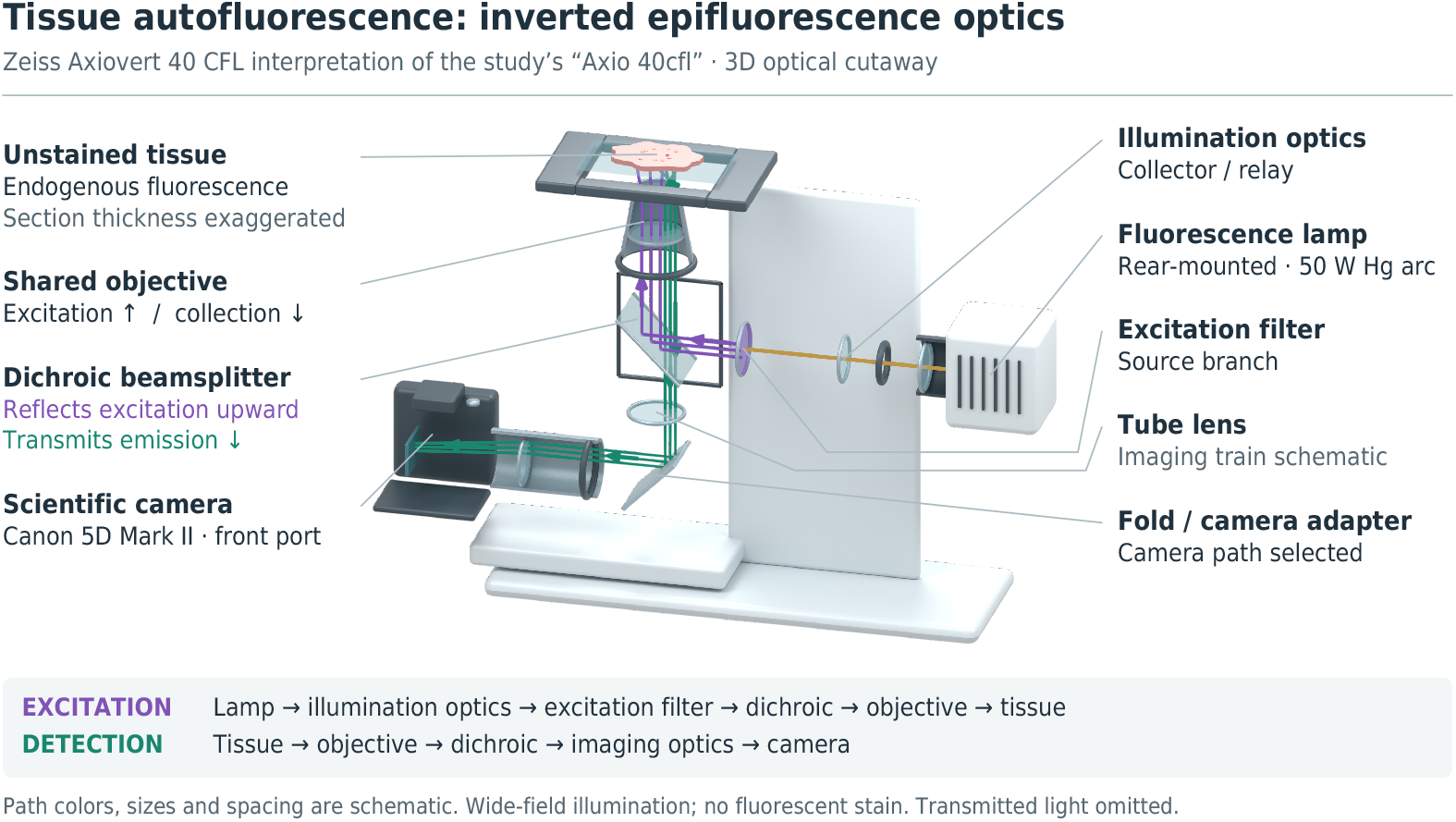
Schematic of tissue epifluorescence imaging. Light from a 50 W mercury arc lamp passes through illumination optics and an excitation filter, reflects from a dichroic beam splitter, and reaches the tissue through a shared objective. Returning fluorescence passes through the dichroic beam splitter and imaging optics to the Canon 5D Mark II camera.

Illumination was provided by a 50 W mercury arc lamp. Color fluorescence images were recorded with a digital camera (Canon 5D Mark II) under ultraviolet (UV) and blue excitation. The DAPI/UV filter configuration used 365 ±12 nm excitation and a 400 nm dichroic beam splitter (BS), with the emission filter removed. The blue (FITC) filter configuration used 480 ±40 nm excitation and a 510 nm dichroic BS, also with the emission filter removed. The exposure time was 1.5 s for both excitations [13]. The spectral response of the system is determined primarily by the filter transmission and the spectral sensitivity of the camera.

### 2.2. Fluorophore concentrations via NMF

As shown in Fig. 3, a 156 *µ*m× 156 *µ*m region was cropped from each fluorescence image. Each cropped image comprised five channels, indexed by *i* according to the excitation and recorded color: UV-Red (*i* = 1), UV-Green (*i* = 2), UV-Blue (*i* = 3), Blue-Red (*i* = 4), and Blue-Green (*i* = 5).

**Fig. 3.**
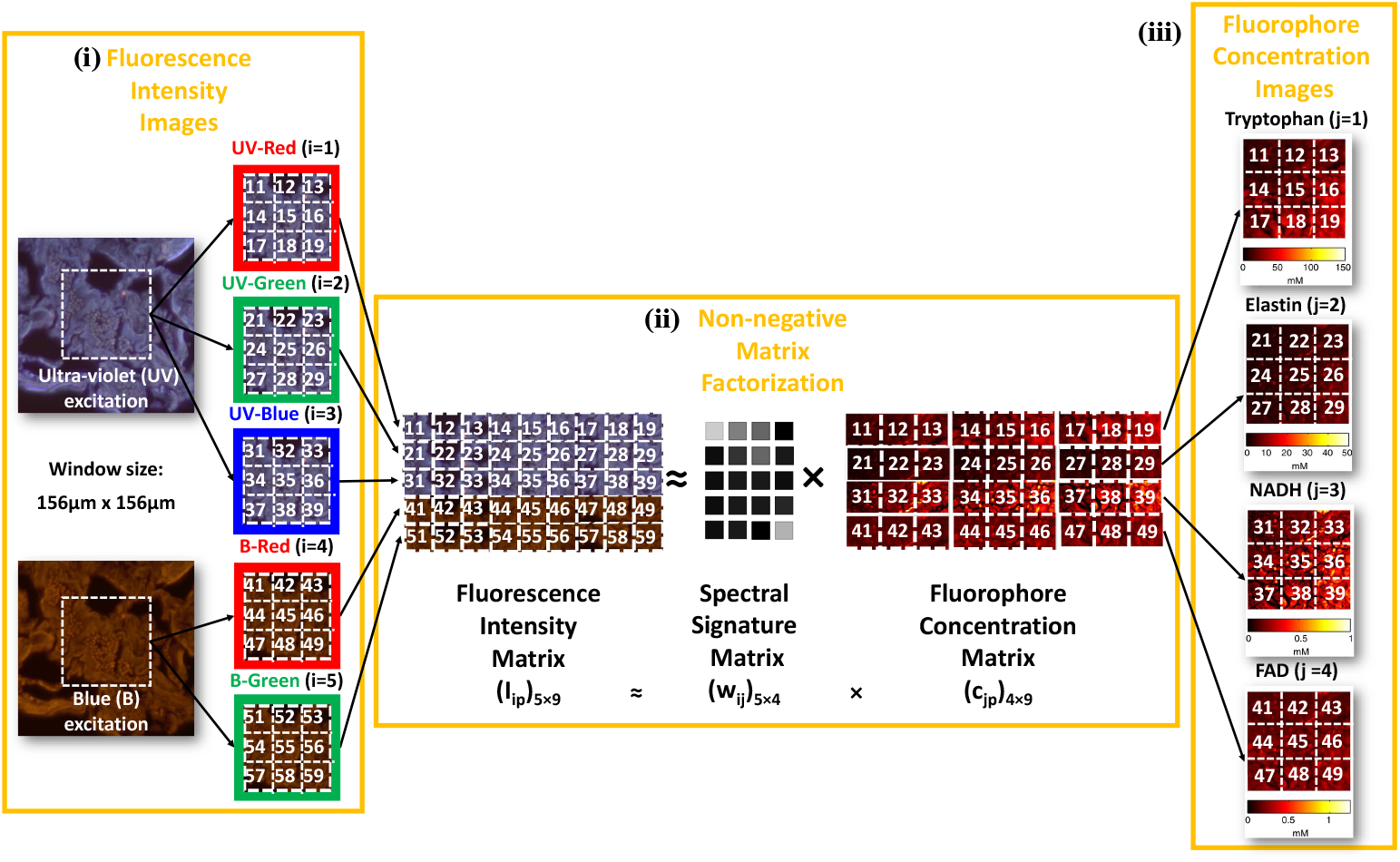
Fluorophore maps obtained from five-channel fluorescence images. (i) Pixel intensities form the measurement matrix. (ii) NMF separates normalized spectral signatures from component fluorescence magnitudes. (iii) Each component row is reshaped into an image; conversion to concentration requires the additional species-specific calibration described in the text.

For each cropped image, the five-channel measurement matrix *I* has dimensions *m* ×*N*, where *m* = 5 and *N* is the total number of pixels. NMF [20] approximates the nonnegative measurements as the product of a normalized spectral-signature matrix *w* and a component-magnitude matrix

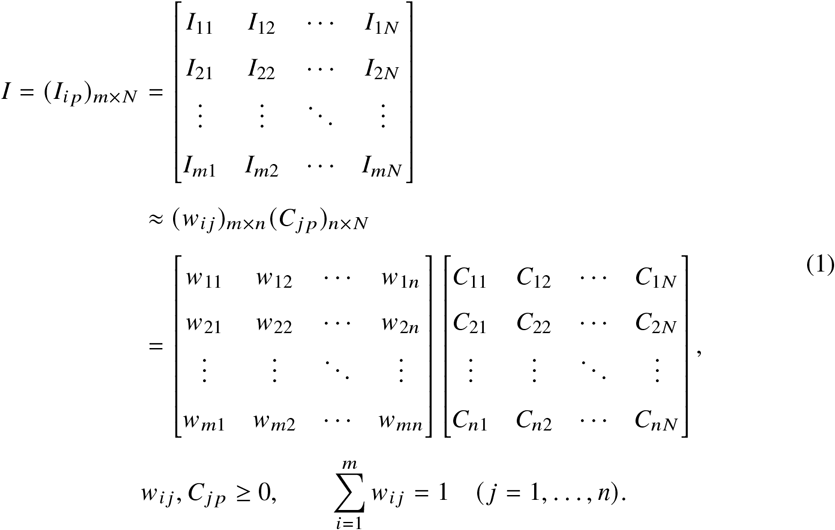

Here *i* = 1, …, *m* indexes measurement channels, *j* = 1, …, *n* indexes the *n* = 4 components (tryptophan, elastin, NADH, and FAD), and *p* = 1, …, *N* indexes pixels. The entry *I*_*i p*_ is the measured intensity, *w*_*i j*_ is the dimensionless fraction of component *j* ‘s fluorescence in channel *i*, and *C* _*j p*_ is its fluorescence magnitude at pixel *p*. The calibrated concentration is *c* _*j p*_ = *C* _*j p*_ /*K* _*j*_, where *K* _*j*_ is the measured fluorescence response per unit concentration under matching acquisition and channel-weighting conditions. Reshaping each row of *c* gives a concentration map in millimolar (mM; Fig. 3). Column normalization alone does not establish molar units.

Column normalization fixes the relative scale of *w* and *C* but does not remove component permutations or guarantee NMF uniqueness. After background removal, the factorization is computed separately for each specimen with the block principal pivoting NMF algorithm, initialized with the spectral signatures of pure fluorophores to reduce this ambiguity. Each extracted component is then identified by matching its spectral signature to a bank of purefluorophore references and calibrated against the reference response measured under the same imaging conditions [13]. When *w* is fixed and has full column rank, five channels constrain four magnitudes per pixel; the channel count alone does not establish the identifiability of a jointly estimated factorization. Spectral overlap and calibration uncertainty remain possible sources of error.

### 2.3. Image dataset and feature extraction

Factorization and calibration produce four concentration maps for each cropped field, one each for tryptophan, elastin, NADH, and FAD. Each map shares the pixel grid of the cropped fluorescence image, and its value at pixel *p* is the calibrated concentration *c* _*j p*_ of fluorophore *j* defined after Eq. (1). A map therefore shows both where a fluorophore is located within the field and how much of it is present at each location [13].

Each map is divided into nonoverlapping sub-images. The same partition is applied to all four maps of a field, so that the features of a sub-image describe one tissue region (Fig. 4). The sub-image is the unit of observation. Subdivision increases the number of observations but not the number of independent specimens or patients.

**Fig. 4.**
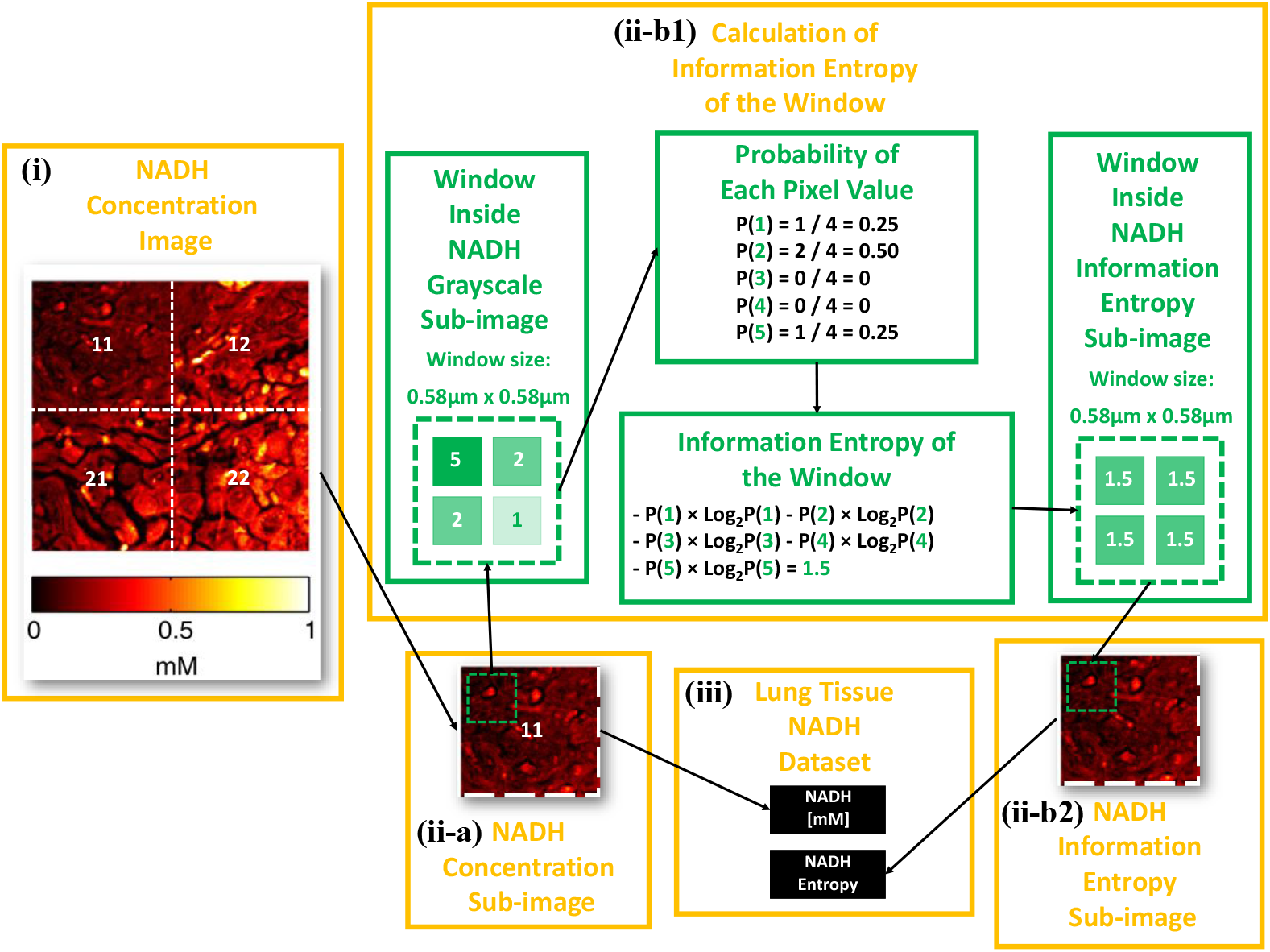
Construction of concentration and information-entropy features, illustrated for NADH. (i) A concentration image is partitioned into sub-images. (ii-a) Pixel concentrations are averaged within each sub-image. (ii-b) Local gray-level histograms in a sliding 9 ×9-pixel window yield an entropy map, which is averaged over the sub-image. (iii) The mean concentration and mean entropy form two features for each fluorophore.

The concentration feature of fluorophore *j* in sub-image *s* is the arithmetic mean of its pixel concentrations,

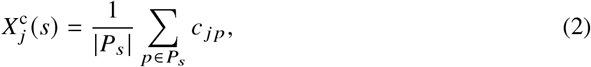

where *P*_*s*_ is the set of pixels in the sub-image and |*P*_*s*_| is their number. This mean describes the abundance of the fluorophore but not how its concentration is distributed within the sub-image.

Information entropy describes that distribution [14]. Each concentration sub-image is first normalized and quantized to 256 gray levels, which assigns an integer level *q* (*p*) {0, …, 255} to each pixel. The probability distribution is then constructed locally. For a 9× 9-pixel window *W*_*r*_ at location *r* (0.58 *µ*m ×0.58 *µ*m), the probability of level *l* is the fraction of the *M* = 81 window pixels at that level,

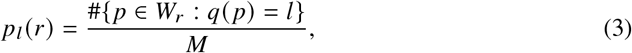

that is, the normalized gray-level histogram of the window. The local Shannon entropy of fluorophore *j*, in bits, is

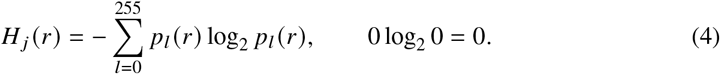

Sliding the window across the sub-image yields a local entropy map. The entropy feature is the arithmetic mean of this map over the set *R*_*s*_ of window locations in the sub-image,

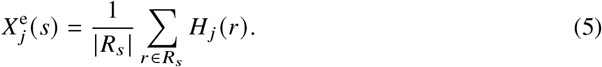

At most 81 levels can be occupied in one window, so *H* _*j*_ (*r*) ranges from 0 bits for a window with a single level to log_2_ 81≈6.34 bits when all 81 pixels occupy different levels.

Entropy depends on the frequencies of the gray levels within a window rather than on their values: adding a constant without clipping or changing bin occupancies leaves entropy unchanged. The window supplies local spatial context, while its histogram discards the arrangement of values within the window. The entropy feature therefore characterizes local intensity diversity but does not uniquely describe tissue architecture.

The features form a table in which each row is one sub-image and each column one variable. Eight predictor columns contain the mean entropy and mean concentration of tryptophan, elastin, NADH, and FAD, and a ninth column, Cancer State, contains the class label. Each subimage is labeled with the diagnosis of the specimen in which it was imaged: *Y* = 0 for normal and *Y* = 1 for cancerous tissue. The label is the probability that the sub-image is cancerous, 0% for a normal and 100% for a cancerous sub-image. The dataset contains 1035 observations (468 normal and 567 cancerous) in nine columns, comprising eight predictors and one target. The same 1035 rows are used for all three feature groups: the concentration-only and entropy-only datasets retain four of the eight predictor columns, and the combined dataset retains all eight. The table does not record the specimen or patient from which each sub-image was taken.

### 2.4. Generalized additive models with structured interactions

A generalized additive model (GAM) represents the conditional mean of a response *Y* given features *X* = ( *X*_1_, …, *X*_*p*_) through an invertible link function *g* [21]:

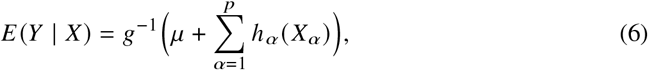

where *µ* is the intercept. Each shape function *h*_α_ depends on one feature, allowing its contribution to be interpreted directly from a curve. A positive main-effect value increases log odds relative to the intercept and other terms; it is not itself a probability or an independently calibrated risk. Pairwise terms extend the model to relationships that cannot be expressed as sums of single-feature contributions. Each function *f*_αβ_ (*X*_α_, *X*_β_) can be visualized as a surface. Restricting interactions to second order provides a practical compromise: higher-order terms can capture additional dependencies, but their contributions are more difficult to interpret directly. GAMI-Net represents each retained curve or surface with a separate neural subnetwork [19]:

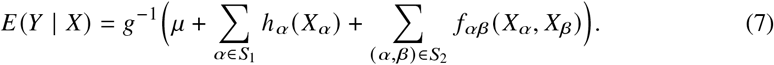

The sets *S*_1_ and *S*_2_ contain the retained main effects and interactions.

For binary classification, *Y* = 1 denotes cancerous and *Y* = 0 normal tissue, so *E* (*Y* | *X*) = *P*(*Y* = 1 | *X*) is the predicted cancer probability. The link *g* is the logit, *g*(π) = log{π/(1 − π)}, and its inverse is the logistic function. In the present dataset, each individual feature *X*_α_ is either the mean concentration 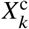 (Eq. (2)) or the mean entropy 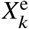 (Eq. (5)) of a fluorophore *k* ∈*K* = {tryptophan, elastin, NADH, FAD}. Each feature pair (*X*_α_, *X*_β_) therefore combines two concentrations, two entropies, or one concentration and one entropy. Grouping the terms of Eq. (7) by these feature types gives the predicted log odds of cancer, η( *X*), and the probabilities of the two classes:

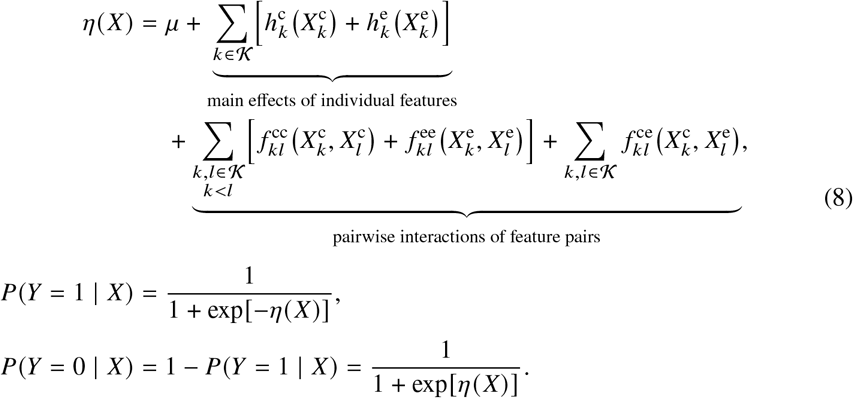

Here 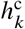 and 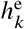 are the main-effect functions *h*_α_ of the concentration and entropy of fluorophore *k*, and 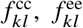, and 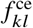 are the interaction functions *f*_αβ_ of concentration–concentration, entropy–entropy, and concentration–entropy pairs; *k* < *l* follows the order in which *K* is listed. A term is zero when its feature or pair is absent from *S*_1_ or *S*_2_. The combined model therefore has eight candidate main effects and 6+ 6 +16 = 28 candidate pairs, including the four concentration–entropy pairs of the same fluorophore (*k* = *l*). The concentration-only model contains only the 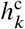 and 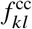 terms, and the entropy-only model contains only the 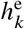 and 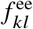 terms.

The main effects and interactions are additive on the log-odds scale. A term with a positive value moves a sub-image toward the cancerous class, and a term with a negative value moves it toward the normal class; the class probabilities depend on the sum η (*X*) rather than on any single term. A sub-image is assigned to the cancerous class when *P* (*Y* = 1 |*X*) ≥τ and to the normal class otherwise. With τ = 0.5, this rule is equivalent to η (*X*) ≥0. ROC AUC summarizes discrimination over all thresholds τ, so the reported results do not depend on a single choice of threshold.

Three constraints in the GAMI-Net framework support interpretation [19]. *Sparsity* removes terms that contribute little to prediction. *Weak heredity* allows an interaction only when at least one parent main effect is retained. *Marginal clarity* penalizes overlap between an interaction and its parent main effects; finite regularization encourages separation rather than guaranteeing exact orthogonality. Centering the term outputs separates their varying contributions from the intercept.

For centered outputs evaluated at the *N*_train_ training observations, importance ratios (IRs) normalize the empirical variances of the fitted terms:

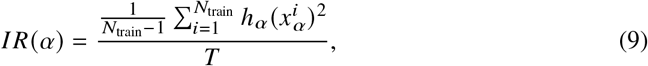

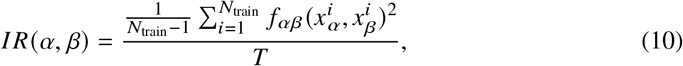

where *T* is the sum of these variances over all retained terms. For *T* > 0, the IRs sum to one. They describe how variation is distributed among model terms over the observed feature distribution and should not be interpreted as causal effects or as fractions of outcome variance explained.

### 2.5. GAMI-Net applied to the fluorophore dataset

The model uses fully connected subnetworks with one input for a main effect and two inputs for an interaction (Fig. 5). The candidate architectures have either two or five hidden layers, each containing 40 nodes. Network depth and activation function are among the hyperparameters considered in Section 2.6. All 28 unique feature pairs are candidates for the eight-feature model; each four-feature model has six possible pairs. The heredity constraint is enabled.

**Fig. 5.**
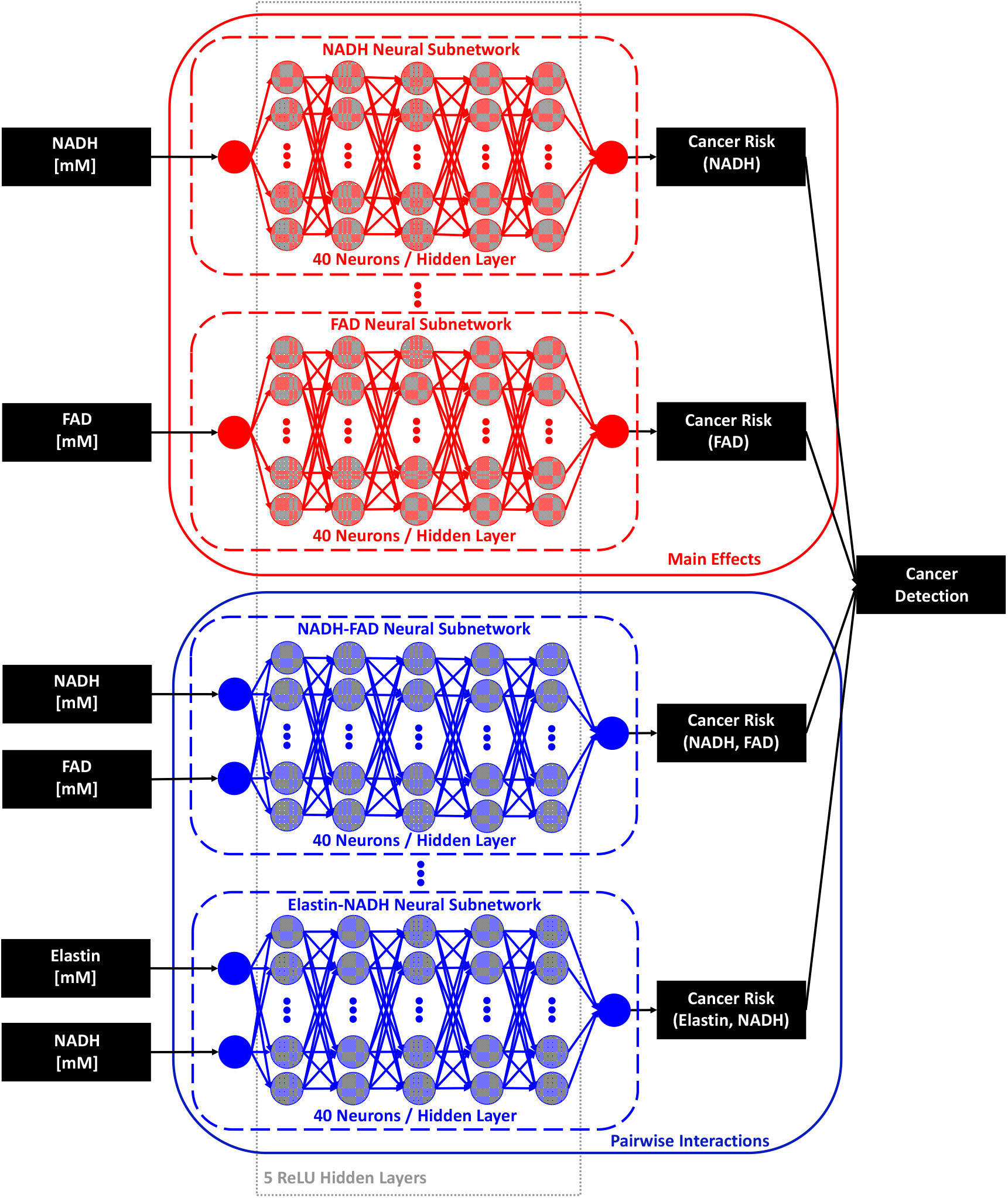
GAMI-Net architecture [19]. A one-input subnetwork estimates each main-effect curve, and a two-input subnetwork estimates each selected interaction surface. Their outputs are summed with an intercept on the log-odds scale. NADH concentration and NADH–FAD illustrate the two types of term. The illustration depicts five ReLU hidden layers; the search also considers two-layer subnetworks.

Training proceeds in three stages: main-effect fitting and selection; interaction fitting and selection with the retained main effects fixed; and joint fine-tuning. Binary cross-entropy loss is minimized using Adam [22].

#### 2.5.1. Cross-validation and data partitions

Each feature group is evaluated by stratified five-fold cross-validation. The 1035 observations are shuffled with a fixed seed (0) and divided into five folds, stratified by the binary cancer-state label so that each fold preserves the normal-to-cancerous ratio of the dataset (468:567). Each fold serves once as the test set, and the remaining four folds form the outer training set. Within each outer training set, a second stratified random split with the same seed assigns 20% of the rows to a validation set and the remaining 80% to a training set. Each predictor is min–max scaled using the training set alone, and the fitted transformation is applied unchanged to the validation and test sets. Training values therefore lie in [0, 1], whereas validation and test values outside the training range can fall outside that interval. Because the partitions depend only on the rows and labels, the three feature groups share identical training, validation, and test rows in each fold and differ only in their predictor columns.

Table 1 gives the exact partition sizes. Every fold has 662 training, 166 validation, and 207 test rows. Each set contains *d* predictor columns, with *d* = 4 for the concentration-only and entropy-only models and *d* = 8 for the combined model, plus the target column `Cancer State`. For the combined model, the training, validation, and test sets of each fold therefore have dimensions 662 ×9, 166× 9, and 207× 9, respectively, and their predictor matrices have dimensions 662× 8, 166 ×8, and 207× 8. The corresponding sets for each four-feature model have dimensions 662× 5, 166 ×5, and 207 ×5. After hyperparameter selection, five normal and five cancerous observations are set aside, and the selected configuration is refitted on the remaining 1025 rows. These rows are partitioned in the same way, giving 656 training, 164 validation, and 205 test rows per fold (Table 1). These ten observations entered the preceding hyperparameter search and therefore do not form an independent holdout set.

**Table 1.** Rows in the training, validation, and test sets of each cross-validation fold, given as total (normal/cancerous). Every set also has *d*+ 1 columns: *d* predictors (*d* = 8 for the combined model and *d* = 4 for each single-group model) and the target Cancer State. The hyperparameter search partitions all 1035 observations; the refit of the selected configuration partitions 1025 observations after ten are set aside.

| Stage | Fold | Training | Validation | Test |
| --- | --- | --- | --- | --- |
| Hyperparameter search | 1–3 | 662 (299/363) | 166 (75/91) | 207 (94/113) |
|  | 4–5 | 662 (300/362) | 166 (75/91) | 207 (93/114) |
| Refit of selected configuration | 1–3 | 656 (296/360) | 164 (74/90) | 205 (93/112) |
|  | 4–5 | 656 (297/359) | 164 (74/90) | 205 (92/113) |

The validation set guides model development. Within each fold, GAMI-Net uses the validation loss for early stopping in each training stage and to determine how many ranked main effects and interactions are retained. Across folds, the mean validation binary cross-entropy selects the hyperparameter configurations. The test set is reserved for evaluating the generalization performance of the selected model. A fold’s test rows are excluded from scaling, fitting, early stopping, and term selection for the model evaluated on them, and only that model’s test-set predictions provide the reported test AUC.

### 2.6. Evaluation unit, model selection, and reporting

The unit of evaluation is a sub-image. The folds in Section 2.5.1 are stratified by cancer state but not grouped by specimen or patient, because the feature table identifies neither. Sub-images from the same specimen can therefore occur in both the training and test sets of a fold, and the reported AUCs measure discrimination under this partitioning rather than performance on independent specimens. Because the specimens were obtained from a cohort of 18 patients [13], grouping by specimen alone would also not guarantee patient independence. Restricting preprocessing to training data addresses a separate source of leakage and does not remove this dependence [23].

A matched feature-group comparison must hold fixed the test-observation identities, cancer-state definition, ROC AUC calculation, preprocessing rules, and tuning procedure. The same filtering and target encoding are applied to all three variants, which also share the same splitting, scaling, and metric rules.

Table 2 describes the exhaustive search: 3 ×3 ×4× 2× 4 = 288 configurations, each evaluated across the five folds. The configuration with the lowest mean validation binary cross-entropy is selected. The selected hyperparameters are held fixed for subsequent fold-wise fitting. Because the selection pools validation losses over all five folds, the test rows of one fold contribute to the training and validation sets of the other folds that inform the choice. This shared selection therefore does not yield a fully nested estimate based on an independent hyperparameter search within every outer training set. For a fold-wise quantity *q* ^(*k*)^, the arithmetic mean across *K* = 5 folds is denoted by 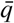. AUCs, importance ratios, and pointwise curve or surface values are reported as “mean ±SD across the five folds.” These SDs describe between-fold dispersion; they are neither confidence intervals nor estimates of uncertainty from independent patient sampling. The five folds share training observations, so fold-level consistency would be descriptive evidence, not five independent replications. No independence-based significance test is applied.

**Table 2.** Hyperparameter values evaluated in the exhaustive grid search.

| Hyperparameter | Tested values | Count |
| --- | --- | --- |
| Mini-batch size | 32, 64, 128 | 3 |
| Activation function | Rectified linear unit (ReLU), hyperbolic tangent, sigmoid | 3 |
| Clarity regularization | $10^{-4}$, $10^{-3}$, $10^{-2}$, $10^{-1}$ | 4 |
| Subnetwork architecture | Two hidden layers of 40 nodes; five hidden layers of 40 nodes | 2 |
| Learning rate | $10^{-4}$, $3 \times 10^{-4}$, $6 \times 10^{-4}$, $10^{-3}$ | 4 |
| Total configurations |  | 288 |

### 2.7. Compute environment

The computational workflow supports training on a central processing unit (CPU) or graphics processing unit (GPU) in a TensorFlow 2.10.0 container.

## 3. Results

### 3.1. Classification performance

Figure 6 reports mean ROC AUCs of 95.9% ±0.7% for concentration-only inputs, 96.4%± 1.6% for entropy-only inputs, and 98.4%± 0.7% for their combination under five-fold sub-image evaluation. These are ROC AUCs, not classification accuracies; the reported uncertainties are SDs across folds.

**Fig. 6.**
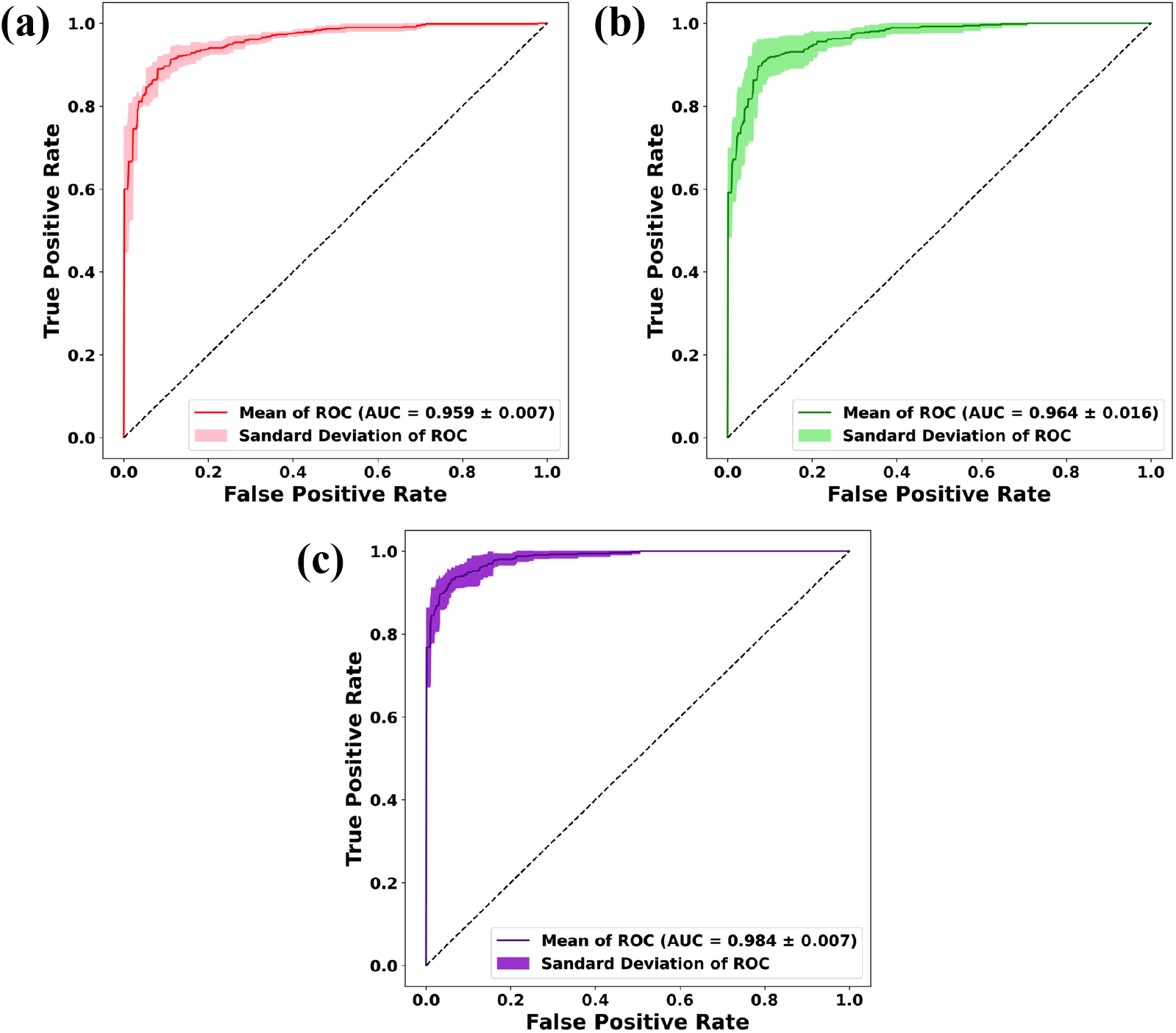
Receiver operating characteristic curves for (a) concentration-only, (b) entropy-only, and (c) combined features. AUCs are reported as mean ±SD across the five sub-image folds, which are not grouped by specimen.

The combined-model mean exceeds the concentration-only and entropy-only means by approximately 2.5 and 2.0 percentage points, respectively. These are differences between reported aggregate AUCs, not relative percentage improvements or averages of paired fold differences. The ordering is consistent with the hypothesis that concentration and entropy provide complementary predictive information. It is not a paired comparison: fold-wise differences between the three models were not evaluated, so it does not show in how many folds the combined model improved, tied, or deteriorated. Neither statistical significance nor generalization to independent specimens follows from the aggregate ordering.

### 3.2. Feature importance

Within each fold, importance is computed from the outputs of the retained terms at that fold’s *N*_train_ training observations. Because each term is centered, the sum of its squared outputs divided by *N*_train_ −1 is its empirical variance on the log-odds scale. Equation (9) divides the variance of the main effect *h*_α_ by *T*, the summed variance of all retained main effects and interactions, and Eq. (10) applies the same normalization to the interaction *f*_αβ_. Each IR is therefore the fraction of the total term variance attributed to the main effect of feature α or to the interaction between features α and β. The IRs within a fold sum to one, and the common factor 1 /(*N*_train_− 1) cancels from each ratio. A term with a large IR produces more variation in the predicted log odds across the observed samples than a term with a small IR. The fold-wise IRs are summarized as mean± SD across the five folds (Fig. 7).

**Fig. 7.**
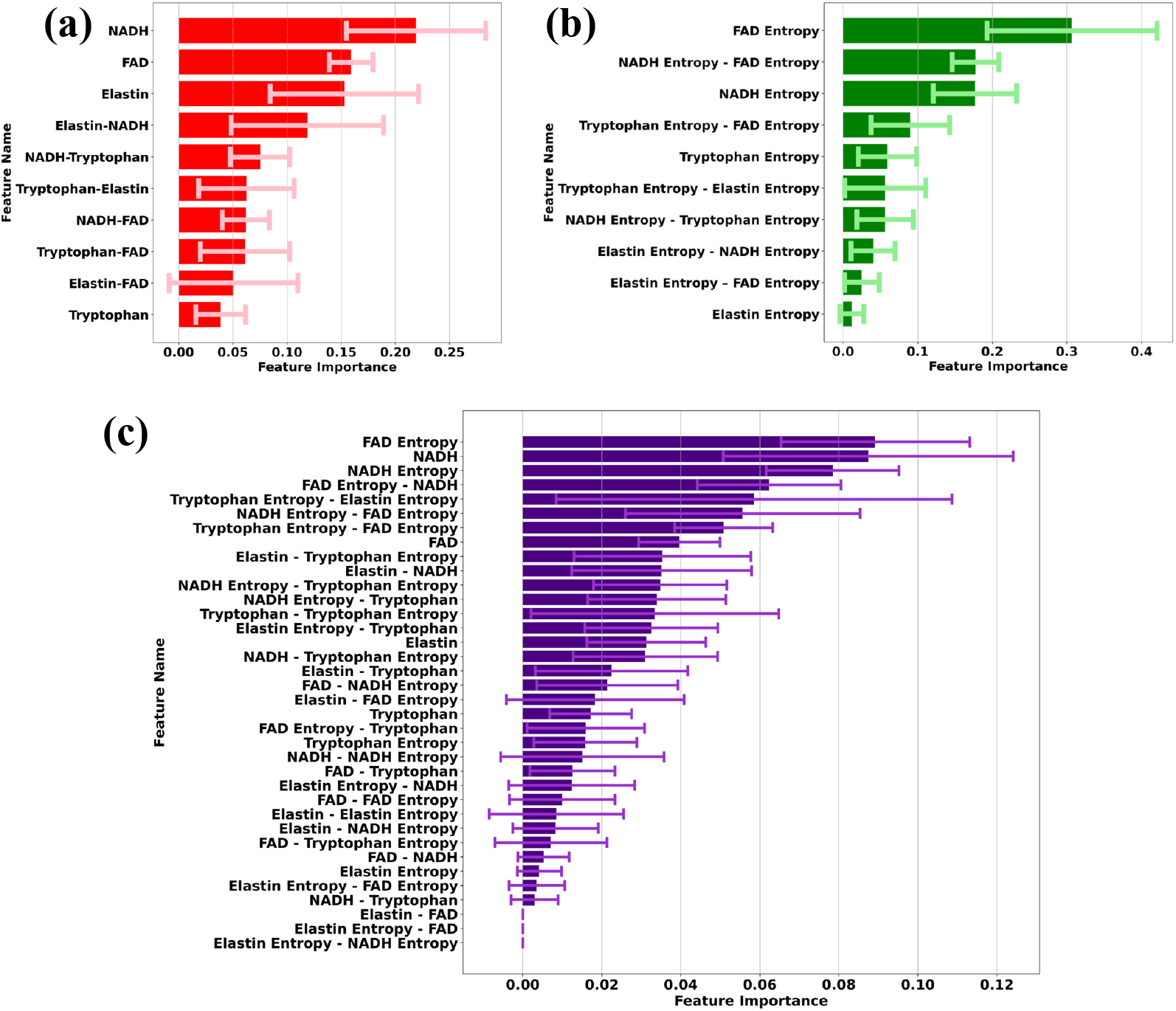
Importance ratios for (a) concentration-only, (b) entropy-only, and (c) combined features, summarized as mean ±SD across the five folds. Importance quantifies variation in the fitted model terms, rather than a causal effect.

In the concentration-only model, NADH, FAD, and elastin concentrations are the three leading main effects, and the elastin–NADH interaction also contributes prominently. In the entropy-only model, FAD entropy and NADH entropy are the leading main effects, and NADH entropy–FAD entropy is the highest-ranked interaction.

In the combined model, FAD entropy, NADH concentration, and NADH entropy are the three leading main effects. FAD entropy–NADH concentration is a prominent interaction, linking a distributional feature with a concentration feature. The three lowest-ranked interactions shown are elastin concentration–FAD concentration, elastin entropy–FAD concentration, and elastin entropy–NADH entropy. Their importance is close to zero, in contrast to that of the dominant main effects. However, these low values alone do not establish that a term was pruned in every fold.

### 3.3. Main-effect cancer-risk curves

Figure 8 compares the main effects shared by the combined model and either the concentration-only or the entropy-only model. The quantity labeled “cancer risk” is an additive contribution to cancer log odds, as defined in Eq. (7). Its sign alone does not determine the predicted class because the intercept, other main effects, and interactions also contribute.

**Fig. 8.**
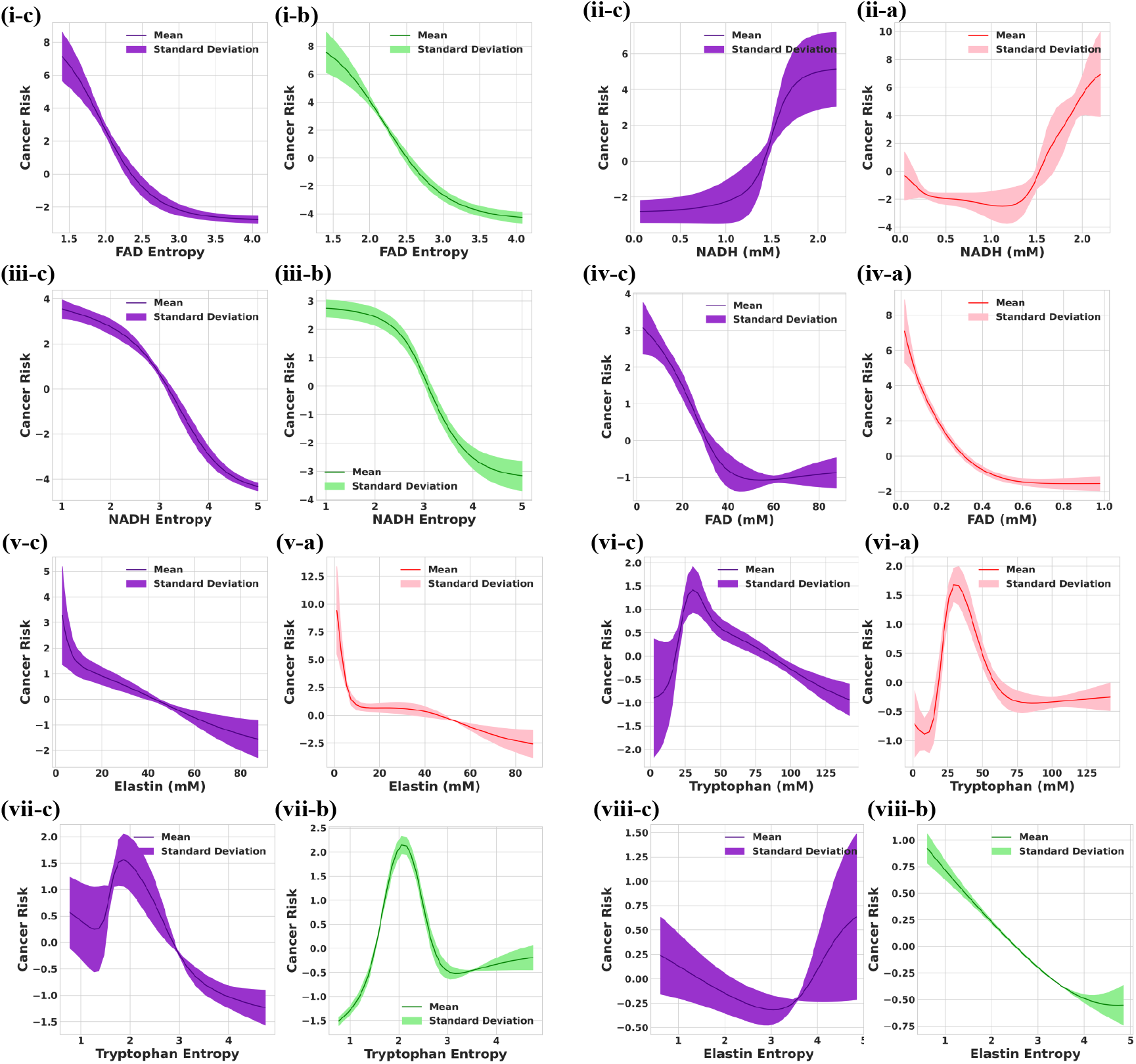
Main-effect contributions to cancer log odds. Each adjacent pair shows the combined model (c, purple) and the corresponding concentration-only (a, red) or entropy-only (b, green) model; bands show the SD across folds. Pairs (i)–(viii) show FAD entropy, NADH concentration, NADH entropy, FAD concentration, elastin concentration, tryptophan concentration, tryptophan entropy, and elastin entropy, respectively.

FAD entropy and NADH entropy show decreasing contributions in both applicable models. Elastin concentration also has a decreasing contribution, and FAD concentration shows a predominantly decreasing response. NADH concentration rises sharply at higher values, although its concentration-only curve is not monotonic throughout the displayed range. Tryptophan concentration and entropy show peaks followed by declining contributions. Elastin entropy is U-shaped in the combined model, whereas its entropy-only contribution is predominantly decreasing. Biological interpretations of these model-dependent associations are considered in Section 4.

### 3.4. Pairwise interaction surfaces

Figure 9 compares two entropy interactions shared by the entropy-only and combined models. Each mean panel shows the interaction term *f*_αβ_ (*X*_α_, *X*_β_) of Eq. (7) as a three-dimensional surface, with the same values projected as filled contours onto the plane of the two features. The color bar beside each pair gives the scale of this contribution to cancer log odds. Positive values raise the log odds and move a sub-image toward the cancerous class; negative values lower them and move it toward the normal class; values near zero leave the prediction determined by the intercept and the other terms. The two models use different color maps: positive contributions are purple and negative contributions orange for the combined model, whereas positive contributions are green and negative contributions purple for the entropy-only model. The combined-model color bars extend to magnitudes of about 8, and the entropy-only color bars to about 4–6, so equal color intensity does not indicate equal magnitude across models.

**Fig. 9.**
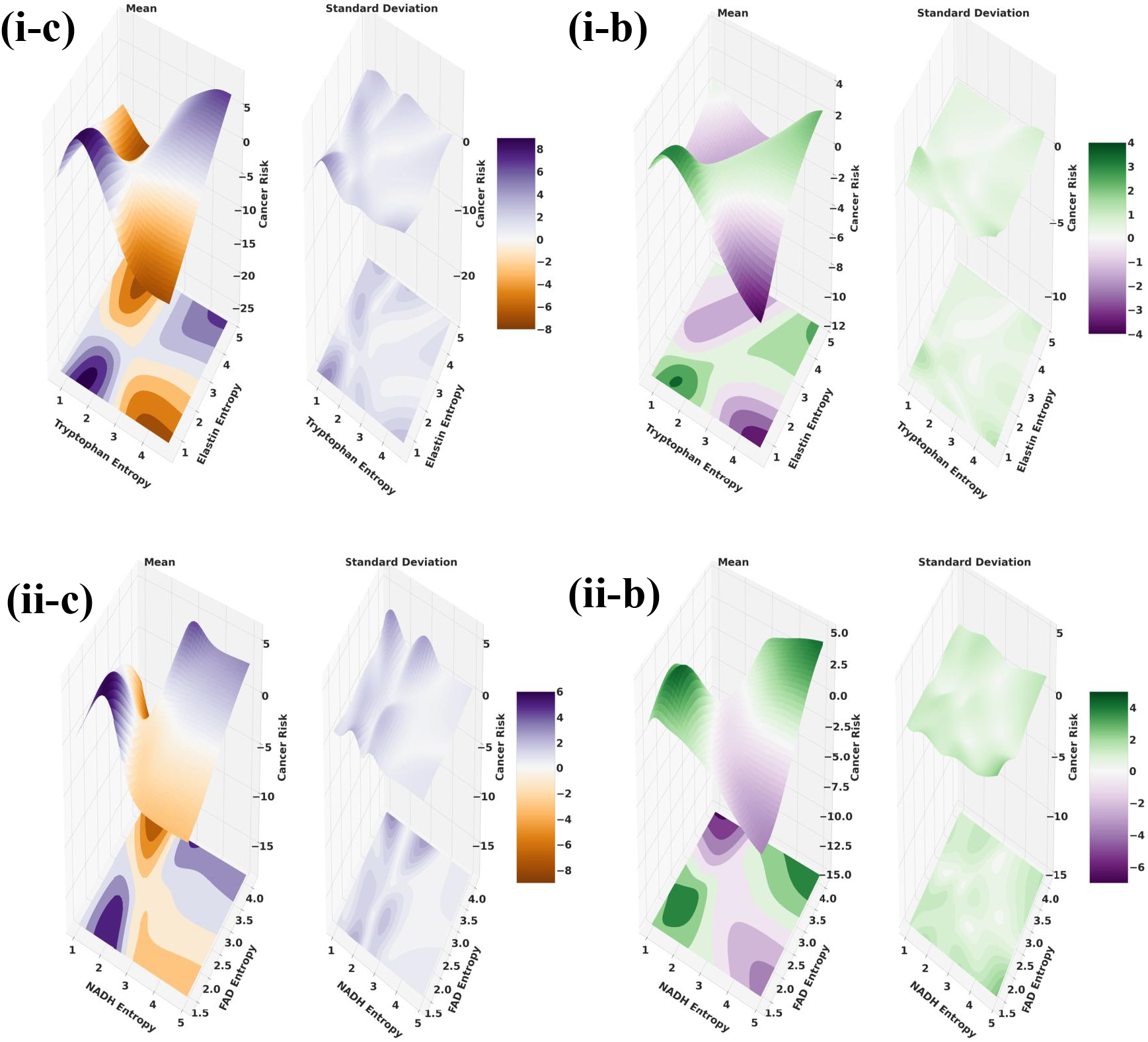
Pairwise contributions to cancer log odds for (i) tryptophan entropy–elastin entropy and (ii) NADH entropy–FAD entropy. Panels (c) and (b) show combined and entropy-only models, respectively. In each mean panel, the contour map on the feature plane projects the surface above it, and the color bar gives the contribution: positive values (purple in c, green in b) raise and negative values (orange in c, purple in b) lower the cancer log odds. These two pairs permit comparison of the same entropy interactions; they do not represent all selected interactions or the two highest-ranked terms in the combined model.

For tryptophan entropy–elastin entropy (Fig. 9(i)), the contribution has a saddle-shaped, checkerboard arrangement in both models. When both entropies are low, near the lower ends of both axes, the contribution is strongly positive and raises the cancer log odds. A second positive region occurs when both entropies are high. In contrast, high tryptophan entropy combined with low elastin entropy gives a negative contribution, which lowers the cancer log odds and moves a sub-image toward the normal class; in the entropy-only model, this is the most negative region of the surface. Low tryptophan entropy combined with high elastin entropy also gives a negative contribution. Contributions are close to zero at intermediate values of both entropies, where the positive and negative regions meet. The effect of tryptophan entropy therefore depends on elastin entropy: at low elastin entropy, increasing tryptophan entropy shifts the contribution from positive to negative, whereas at high elastin entropy it shifts the contribution from negative to positive.

NADH entropy–FAD entropy (Fig. 9(ii)) shows the same concordant–discordant arrangement. Low NADH entropy with low FAD entropy yields a strongly positive contribution in both models, and high NADH entropy with high FAD entropy yields a second positive region. The discordant combinations are negative: the most negative contributions occur at low-to-intermediate NADH entropy combined with high FAD entropy, while high NADH entropy combined with low FAD entropy gives a moderately negative contribution. In the combined model, a band of slightly negative values near the middle of the NADH-entropy axis separates the low-NADH-entropy and high-NADH-entropy regions. As for the first pair, the sign of the contribution associated with one entropy reverses with the level of the other.

For both pairs, the two models agree in the locations and signs of the main positive and negative regions, which indicates that this interaction pattern is not specific to one feature set. Agreement in arrangement does not imply equal magnitude, because the color scales differ. An individual sub-image’s predicted class cannot be read from these maps alone: the interaction value is added to the intercept, the main effects in Fig. 8, and any other retained interactions to give η (*X*) in Eq. (8). Because marginal clarity encourages separation of interactions from their parent main effects, the surfaces approximately describe departures from the sum of those main effects on the log-odds scale; they do not represent probabilities on their own. The surfaces are drawn over the whole displayed grid and do not show where observations are dense or sparse, so the values at the corners of the feature ranges should be interpreted with caution. A predictive interaction does not by itself establish a biological interaction or mechanism. In particular, an interaction between NADH and FAD entropies is distinct from the concentration or fluorescence ratio used to assess redox state.

## 4. Discussion

The reported combined-feature mean ROC AUC of 98.4% ± 0.7% exceeds the concentration-only and entropy-only means by approximately 2.5 and 2.0 percentage points, respectively. This aggregate ordering supports the hypothesis descriptively: it is consistent with an advantage from combining abundance and local distributional information.

The interpretable decomposition identifies both feature groups among the leading contributors. FAD entropy, NADH concentration, and NADH entropy dominate the combined model’s main effects, while FAD entropy–NADH concentration is a prominent pairwise term. The concentration-only model also uses the elastin–NADH interaction, and NADH entropy–FAD entropy leads the entropy-only interactions. Importance measures variation in fitted terms over the observations; it neither quantifies causal influence nor proves that a feature supplies unique information independent of correlated inputs. Likewise, a model interaction expresses a predictive departure from additivity, not a demonstrated biochemical interaction.

The learned contributions do not increase uniformly with concentration or entropy. FAD- and NADH-entropy contributions decrease in both applicable models, tryptophan contributions are nonmonotonic, and elastin entropy is U-shaped in the combined model but predominantly decreasing in the entropy-only model. These differences show why the functional forms matter beyond a ranking of features. They also rule out interpreting the classifier through a universal claim that greater image entropy indicates cancer.

The shape functions make the model’s use of biochemical and distributional information explicit. Decreasing contributions with FAD concentration, together with higher contributions at elevated NADH concentration, are qualitatively compatible with a lower FAD-based optical redox ratio in glycolytic precancerous tissue models [4]. However, separate additive terms are not a redox-ratio measurement, and a conditional NADH curve need not follow the unadjusted concentration difference between tissue classes. Fixed-tissue fluorescence, binding state, and tissue composition further limit a direct interpretation in terms of metabolic flux.

The U-shaped elastin-entropy contribution in the combined model suggests that both low and high local intensity diversity can be associated with elevated model scores. One possible explanation is that relatively uniform, elastin-depleted regions and heterogeneous regions of matrix remodeling represent different tissue states associated with malignancy. Extracellular-matrix remodeling provides a biological basis for investigating this possibility [9], but the model does not identify the underlying tissue compartments. The predominantly decreasing entropy-only curve also shows that the inferred relationship depends on the other features included in the model. Histological colocalization would be needed to test this interpretation.

The peaks in the tryptophan concentration and entropy curves may likewise reflect a combination of tissue composition, protein-associated fluorescence, and metabolic states. Tryptophan catabolism can support tumor immune escape [11, 12], but fluorescence does not separately quantify free tryptophan, its metabolites, or immune activity. An increase followed by a decrease therefore suggests a nonlinear predictive association rather than a direct dose response. Correlated inputs and sparse observations near the extremes of the feature ranges can also shape these curves. The wide uncertainty bands in some regions argue against assigning precise biological thresholds. For context, Xu et al. (2017) reported SVM classification accuracies of 99.8% using absolute concentrations and 93.9% using relative fluorescence strengths [13]. Their noncancer group included perilesional and normal tissue, whereas the present task compares normal and cancerous tissue. Accuracy depends on a classification threshold, whereas ROC AUC measures ranking across thresholds. The present AUC therefore cannot be interpreted as an accuracy reduction relative to their 99.8%, and differences in observations, feature construction, and validation protocol preclude a direct benchmark comparison. Establishing an accuracy–interpretability tradeoff would require retraining both models with the same cohort, features, partitions, and metrics.

Several limitations define the scope of these findings. The analysis uses 36 normal and 52 cancerous specimens from a single BioMax tissue-microarray cohort, with no external validation. Subdivision into sub-images introduces dependent observations, and ungrouped folds can yield optimistic estimates when specimen-specific properties appear in both training and test data. Validation grouped by specimen and, where applicable, by patient is needed, followed by evaluation in an independent cohort. Hyperparameter selection should also be repeated within each outer training fold to obtain a fully nested performance estimate. Fold SDs do not quantify these sources of optimism.

The eight constructed features summarize concentration and local entropy rather than representing the images directly. The local histogram omits arrangement within each neighborhood, so the entropy features cannot identify tissue architecture or cellular compartments uniquely. They may discard diagnostically relevant morphology, and their values depend on image sub-division, grayscale quantization, and calibration. NMF normalization, spectral matching, and calibration provide an interpretable representation of the measurements but do not guarantee unique separation or eliminate specimen-dependent spectral error. Finally, model contributions are observational associations: neither high importance nor a prominent interaction establishes a causal cancer mechanism.

## 5. Conclusion

The reported aggregate ROC AUCs favor combining fluorophore concentration and local histogram entropy over either feature group alone for normal-versus-cancerous lung-tissue classification: the combined-model mean is 98.4%±0.7%, approximately 2.5 and 2.0 percentage points above the concentration-only and entropy-only means. GAMI-Net exposes the main effects and pairwise functions underlying predictions on the log-odds scale, providing an interpretable account of nonlinear predictive associations.

The decomposition shows how the two descriptions of the same fluorophore maps contribute. Concentration and entropy features both rank among the leading terms: FAD entropy, NADH concentration, and NADH entropy dominate the combined model’s main effects, and the prominent FAD entropy–NADH concentration interaction joins a distributional feature with an abundance feature. Local entropy therefore enters predictions alongside concentration rather than duplicating it, consistent with the premise that sub-images with the same mean concentration can differ in the distribution of their values. The learned relationships are not uniform: NADH- and FAD-entropy contributions decrease, tryptophan contributions are nonmonotonic, and elastin entropy is U-shaped in the combined model. The tryptophan entropy–elastin entropy and NADH entropy–FAD entropy surfaces share a concordant–discordant arrangement in both the entropy-only and combined models, in which the contribution associated with one entropy reverses sign with the level of the other. A ranking of features alone would not reveal these forms, and they do not support a universal association between greater image entropy and cancer.

The significance of this work lies in coupling label-free measurement with an explicit model decomposition. CFM derives fluorophore maps from unstained tissue using endogenous contrast, and GAMI-Net expresses its predictions as a sum of curves and surfaces over physically defined features: fluorophore concentrations and local entropies in bits. Each learned contribution can therefore be inspected, compared between feature sets, and related to tissue biochemistry and organization. Several contributions are qualitatively compatible with known cancer-associated changes, including the opposing NADH- and FAD-concentration contributions with a shift toward glycolytic metabolism and the elastin-entropy contribution with extracellular-matrix remodeling. These correspondences do not establish mechanisms; instead, they turn an accurate classifier into specific hypotheses that can be tested, for example by histological colocalization of elastin entropy with matrix compartments or by measurements that distinguish free from protein-bound cofactors. Comparing the same term across models also identifies relationships that depend on the other features included, as the differing elastin-entropy curves illustrate. The same feature construction and model can in principle be applied to other endogenous fluorophores and tissue types imaged by CFM.

These conclusions apply to sub-image evaluation within a single tissue-microarray cohort. Because the folds were not grouped by specimen and hyperparameters were selected across folds, the reported AUCs do not estimate performance on new specimens or patients. Paired fold-wise comparison of the three feature groups, specimen- and patient-grouped nested validation, confirmation of the concentration calibration, and evaluation in an independent cohort are required before the combination of concentration and entropy can be assessed as a diagnostic aid for unstained lung tissue.

